# Nitrogen utilisation by the metabolic generalist pathogen *Mycobacterium tuberculosis*

**DOI:** 10.1101/385278

**Authors:** Aleksandra Agapova, Debbie M. Hunt, Michael Petridis, Acely Garza-Garcia, Charles D. Sohaskey, Luiz Pedro S. de Carvalho

## Abstract

Bacterial metabolism is fundamental to pathogenesis and has a dominant effect on bacterial killing by antibiotics. Here we explore how *Mycobacterium tuberculosis* utilises amino acids as nitrogen sources, using a combination of bacterial physiology and stable isotope tracing coupled to liquid chromatography mass spectrometry metabolomics methods. Our results define core properties of the nitrogen metabolic network from *M. tuberculosis*, such as: (i) the lack of homeostatic control of amino acid pool sizes; (ii) similar rates of utilisation of different amino acids as sole nitrogen sources; (iii) improved nitrogen utilisation from amino acids compared to ammonium; and (iv) co-metabolism of nitrogen sources. Finally, we discover that alanine dehydrogenase, is involved in ammonium assimilation in *M. tuberculosis*, in addition to its essential role in alanine utilisation. This study represents the first in-depth analysis of nitrogen source utilisation by metabolic generatlist *M. tuberculosis* and reveals a flexible metabolic network with characteristics that are likely product of evolution in the human host.

---

Tuberculosis, caused by the bacillus *Mycobacterium tuberculosis*, is now the greatest cause of death by a single infectious agent, surpassing deaths caused by HIV/AIDS^1^. Failures in drug discovery programmes aimed at finding transformative antitubercular agents are thought to partially derive from by our incomplete understanding of bacterial phenotypic diversity in the host ^2^. Bacterial metabolic flexibility is thought to be essential for survival in a variety of niches, where low pH, low oxygen tension, presence of reactive oxygen species, and scarcity of nutrients are commonly found.

In contrast to the wealth of knowledge on carbon metabolism^3,4^, very little is known about nitrogen metabolism, in particular, we do not understand what are the essential features of nitrogen metabolism in *M. tuberculosis*^5^. For example, while we understand how post-translational regulation of nitrogen metabolism operates in mycobacteria^6–13^, transcriptional regulation of nitrogen metabolism in *M. tuberculosis* is largely unknown (**Fig. S1**). The transcriptional factor *GlnR* does not perform canonical functions in mycobacteria^14,15^. Nitrogen utilisation by different mycobacterial species appears to be very different, not only due to inherent growth rate differences but also lag phase and final biomass achieved (**Fig. S2**). Importantly, the vast majority of studies to date focused exclusively on either ammonium (NH_4_^+^) as the sole physiologically relevant nitrogen source^14,16^ or employed surrogate fast-growing species such a *M. smegmatis* instead of *M. tuberculosis^16^*. In 2013-14 two key studies unveiled an important aspect of host-relevant nitrogen metabolism in *M. tuberculosis*, namely that host amino acids such as L-aspartate (Asp) and L-asparagine (Asn) were shown to be important sources of nitrogen during infection^17,18^. These findings open a new avenue in host-*M. tuberculosis* relevant metabolism, revealing the use of organic nitrogen sources by *M. tuberculosis* during infection. In addition, there is now overwhelming evidence on the importance of amino acids during infection, highlighted by profound infection attenuation observed with genetic knockout strains^19,20^. Here we describe our studies aimed at exploring core nitrogen metabolic network of *M. tuberculosis* (**Fig. 1**), using amino acids as nitrogen sources.

**Figure 1.**
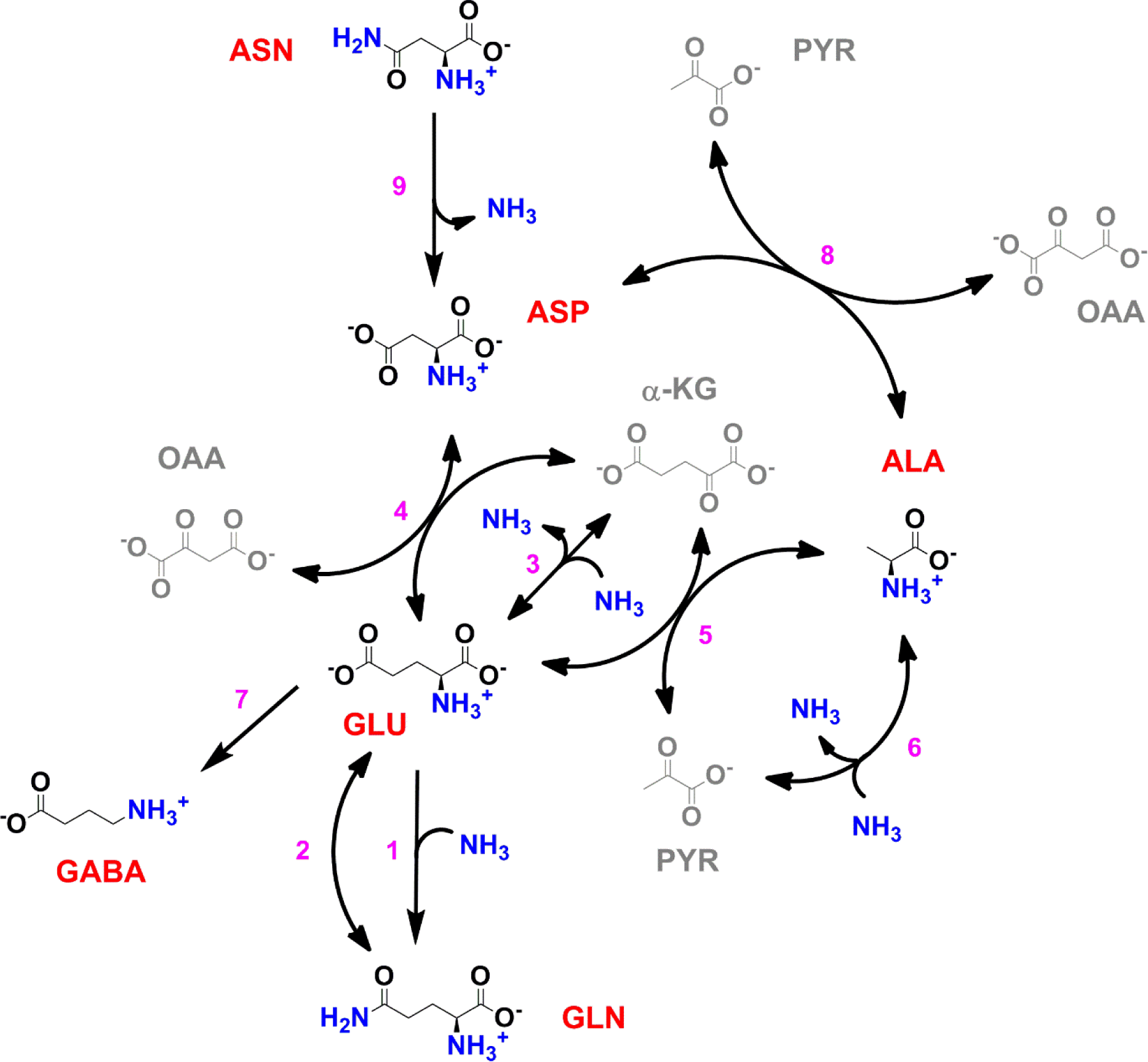
Scheme of the core nitrogen metabolic network of *M. tuberculosis*. 1 – Glutamine synthetase (*glnA*1); 2 – glutamate synthase (*gltBD*); 3 – glutamate dehydrogenase (*gdh*); 4 – glutamate/oxaloacetate transaminase (*aspB*); 5 – glutamate/pyruvate transaminase (*aspC*); 6 – alanine dehydrogenase (*ald*); 7 – glutamate decarboxylase (*gadB*); 8 – aspartate/pyruvate transaminase (*aspC*); 9 – asparaginase (*ansA*).

## RESULTS

### *M. tuberculosis* can take up all proteinogenic amino acids

To investigate amino acid uptake and utilization by *M. tuberculosis*, we transferred bacteria-laden filters after 5 days’ growth on 7H10 media to individual fresh 7H10 agar plates containing 1 mM of each of the 20 proteinogenic amino acids. Cells were harvested 17 hours post-shift. Metabolites were extracted, separated, identified and quantified by liquid-chromatography high-resolution mass spectrometry, following procedures described elsewhere^21,22^. The majority of amino acid intracellular pool sizes vary only modestly when *M. tuberculosis* is grown in media containing sole nitrogen sources different to NH_4_Cl (**Fig. 2a, b**). However, an increase in intracellular pool size is observed for Gly, Ala, Val, Ile, Met, Pro, Phe, Tyr, Trp, Ser, Thr, Arg, and His when *M. tuberculosis* was cultured with the cognate amino acid as sole nitrogen source (highlighted in the diagonal of **Fig. 2a**). Importantly, all amino acids present as sole nitrogen source alter the pool size of the cognate amino acid and/or other amino acid in *M. tuberculosis*, demonstrating that they are taken up. On **Fig. 2b**, the data from **Fig. 2a** is replotted to illustrate individual amino acids changes (final concentrations in samples) obtained with *M. tuberculosis* grown with different amino acids as sole nitrogen sources. With few exceptions (Met and Trp) no change is observed in the summed amino acid pool size, when *M. tuberculosis* is incubated with different amino acids as the sole nitrogen source. **Fig. 2c** contains the data from **Fig. 2b** replotted as summed fold-change, compared to NH_4_Cl. Trp and His as sole nitrogen sources display significant effects on the summed abundance, but most of the other amino acids do not significantly affect overall pool size. In other words, Trp and His are readily taken up by Mtb and stored at high concentrations. **Fig. 2d** shows data for all amino acids, independent of nitrogen source. It is apparent that most amino acid concentrations are not significantly altered in different nitrogen sources, while the concentrations of Pro, Asp, Gln, Glu and Ala vary depending on the sole nitrogen source used. **Fig. 2e** illustrates amino acid levels observed when the cognate amino acid or NH_4_Cl were used as sole nitrogen source. Nearly all amino acid pool sizes are altered when the cognate amino acid is present in the growth medium, as sole nitrogen source. This data is also highlighted on the diagonal in **Fig. 2a**. Curiously, no change is observed in Leu, Asn, Gln, Asp, Glu and Lys when the respective cognate amino acid was added to the growth medium. **Fig. 2f** illustrates the modest changes in the concentration of Gln in *M. tuberculosis* when different amino acids or NH_4_Cl are present in the growth medium. This result suggests that Gln might not be the indicator of nitrogen levels in mycobacteria.

**Figure 2.**
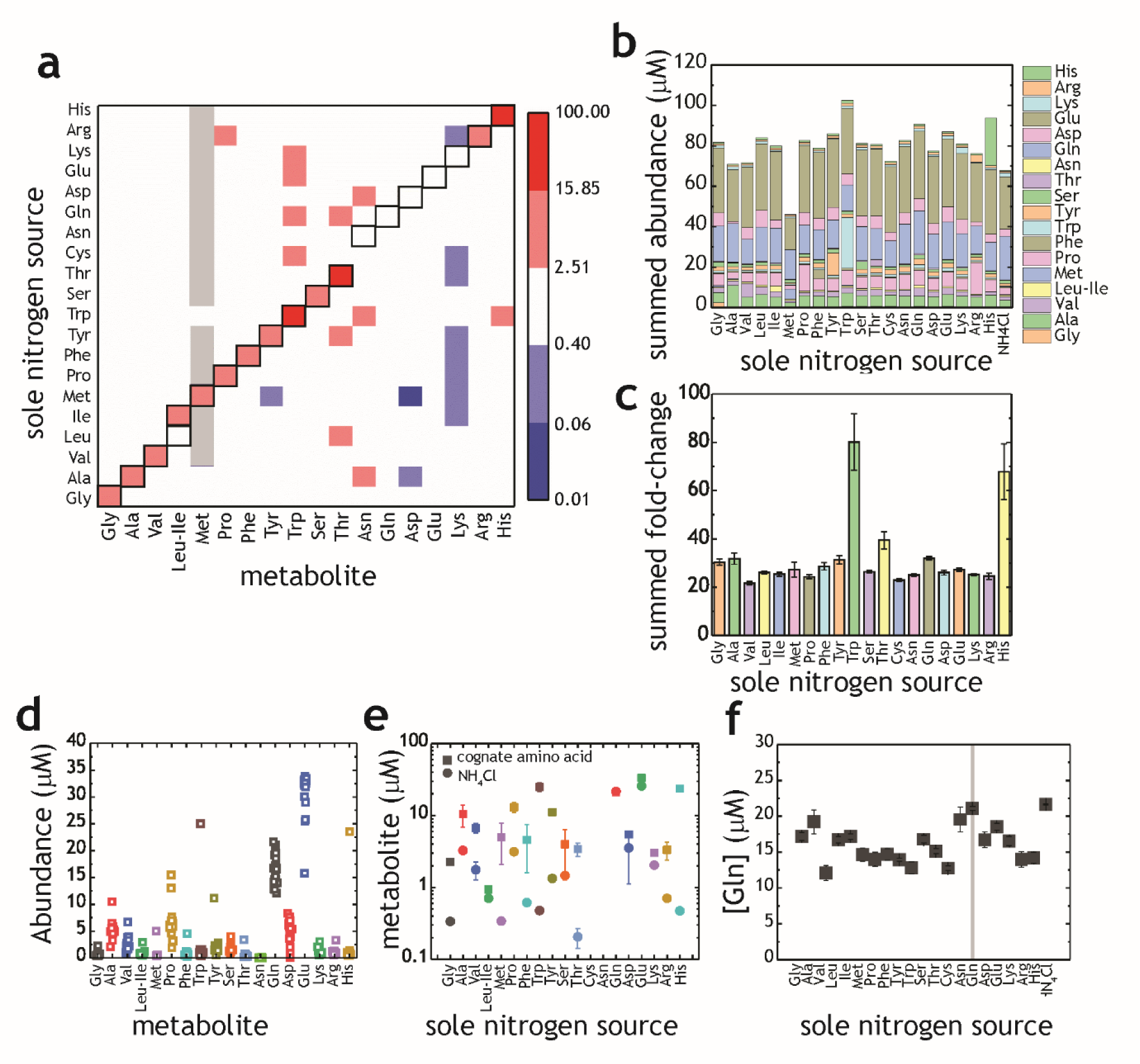
Proteinogenic amino acids as sole nitrogen source for *M. tuberculosis*. (**a**) Heatmap illustrating the changes in amino acids (X-axis) when *M. tuberculosis* is grown on each individual amino acid as sole nitrogen source (Y-axis). Data shown as fold-change (aa/NH_4_^+^). Grey squares indicate that abundance of a particular metabolite was too low to be quantified. Cysteine was undetectable in all conditions and was omitted from this plot. Panels (**b-f**) are re-plots of the data shown in panel (**a**). (**b**) Summed abundance of amino acids in each amino acid as sole nitrogen source. (**c**) Data from panel (**b**) presented as fold-change over NH_4_^+^. (**d**) Amino acid abundances irrespective of the sole nitrogen source used, highlighting the variation on each amino acid in different nitrogen sources. Each symbol represents the average concentration obtained with a single individual nitrogen source. (**e**) Amino acid concentrations in NH_4_^+^ and in medium containing the cognate amino acid as sole nitrogen source. (**f**) Concentration of Gln in extracts from *M. tuberculosis* grown on different amino acids as sole nitrogen source. All concentrations are final concentrations in lysates obtained from approximately 10^9^ cells, and not concentrations per cell. Data is the average of three biological replicates and representative of two independent experiments.

Overall, these results indicate that *M. tuberculosis* does not control the pool sizes of most amino acids homeostatically, *i.e.* intracellular concentrations rise or fall depending on extracellular amino acid/nitrogen source availability.

### Amino acids are superior nitrogen sources, compared to NH_4_^+^

Before carrying out an in-depth analysis of nitrogen metabolism we investigated whether or not the medium used to culture *M. tuberculosis* prior to switching to media with defined sole nitrogen sources could lead to false results. Pre-culture media composition has been shown to affect carbon metabolism^21^. We ‘pre-cultured’ *M. tuberculosis* in either standard Middlebrook 7H9 broth (containing Glu and NH_4_^+^) or a 7H9^NH4+^ broth (a synthetic version of Middlebrook 7H9 broth, with NH_4_^+^ as sole nitrogen source), prior to the experiment in 7H9^NH4+^ broth. When pre-conditioned in standard 7H9, growth of *M. tuberculosis* in 7H9^NH4+^only broth led to a significantly higher biomass accumulation than when pre-conditioned in 7H9^NH4+^ broth (**Fig. S3**). Therefore, pre-adaptation in the nitrogen source that will be tested is essential to avoid overestimation of growth, particularly in poor nitrogen sources. Hence, all experiments were carried out with cultures that were pre-adapted in media of identical composition to the test media for at least 3 days (unless otherwise stated).

**Fig. 3a** illustrates representative growth curves obtained in Glu, Gln, Asp, Asn and NH_4_Cl, as sole nitrogen sources. All four amino acids were superior nitrogen sources to NH_4_Cl, at all concentrations tested (**Fig. 3b and c**), both in terms of doubling rate and final biomass generated. It is noteworthy that pre-adaptation in medium with NH_4_Cl as sole nitrogen source, reveals that *M. tuberculosis* can only optimally utilise NH_4_^+^ as sole nitrogen source until up to 0.25 g/L (4.67 mM). Higher NH_4_^+^ concentrations appeared to be toxic. Based on final biomass produced the following order represents the preferential utilisation of sole nitrogen sources: Glu > Asp > Asn > Gln > NH_4_^+^. When pre-adapted cultures were grown in medium containing no nitrogen source, growth persisted in all cases (**Fig. 3d**). This limited growth is likely due to the low levels of ferric ammonium citrate (0.04 g/L) added to the medium as an iron source. Interestingly, not all cultures grew identically. Growth was different depending on nitrogen source: cells pre-adapted to Asp, Asn and NH_4_Cl allowed growth to an OD ~1.0, while those pre-adapted to Glu and Gln grew to OD ~0.2. To confirm the ‘metabolic conditioning effect’ induced by the pre-adaptation medium, we sub-cultured cells after 15 days into fresh medium. Pre-adaptation with Glu and Gln again led to poor growth, while cells derived from medium containing Asp, Asn and NH_4_Cl grew to an OD ~ 1, in a concentration-dependent manner (**Fig. 3e**).

**Figure 3.**
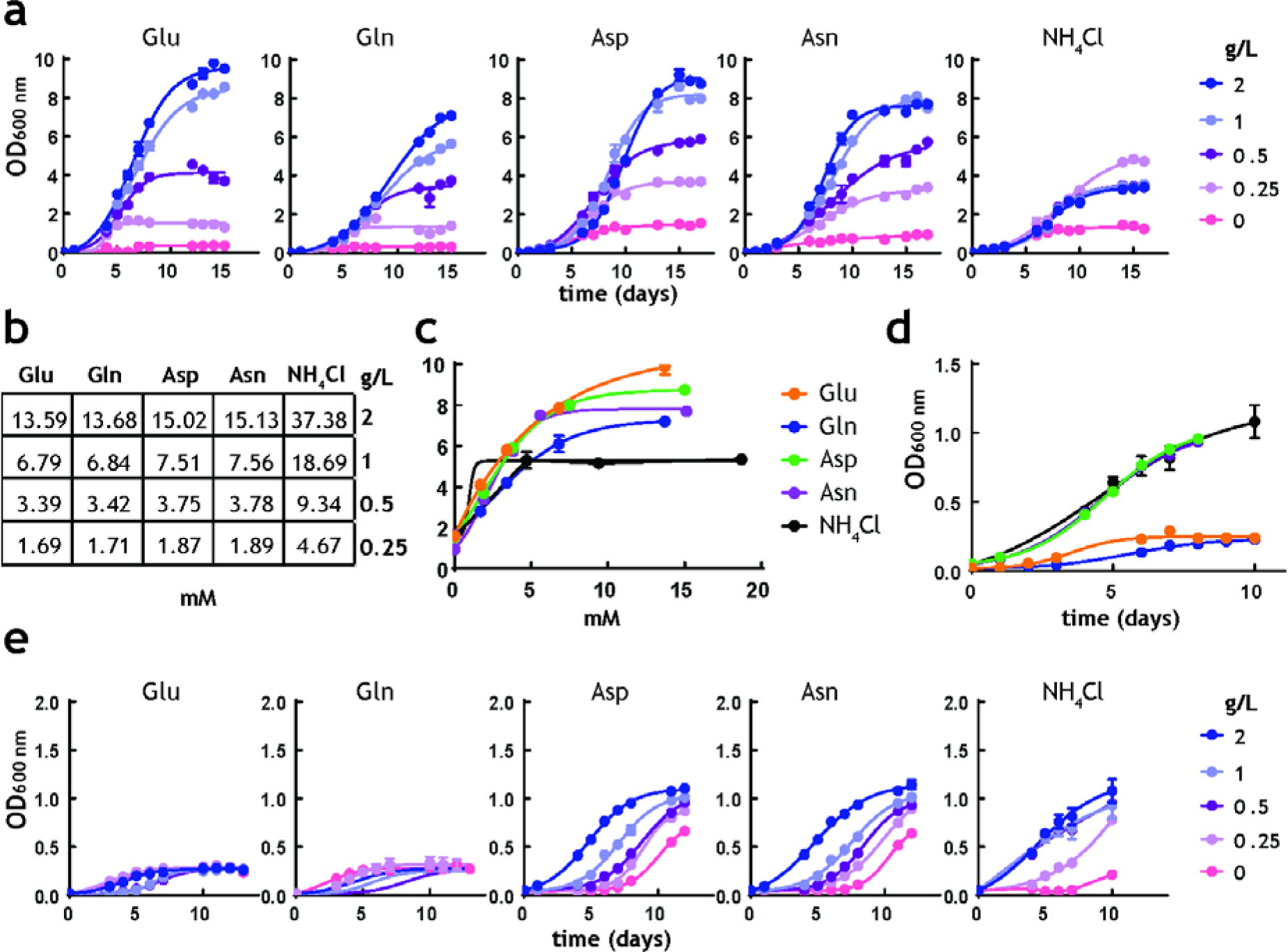
Analysis of *M. tuberculosis* growth in pre-adapted nitrogen cultures. (**a**) Growth curves in 7H9Nx broth (sole nitrogen source). (**b**) Table with g/L to mM conversions for each nitrogen source used. (**c**) Replot of final biomass achieved for each nitrogen source, after 15 days (**a**). Solid lines are the fit to a hyperbolic equation, describing saturation. (**d**) Re-plot of data at no-nitrogen from **a**, illustrating different residual growth. (**e**) Growth curves in media lacking nitrogen, after cultures were grown for 15 days on nitrogen media (**a**), illustrating differential storage of different amino acids. Symbols are data and solid lines in growth curves are the fit to a sigmoidal equation describing bacterial growth. Data are representative of two independent experiments. Error bars are standard error of the mean.

Taken together, these results reveal that the amino acids Glu, Gln, Asp, and Asn are superior to NH_4_^+^ as sole nitrogen sources for *M. tuberculosis*, leading to high biomass and faster growth.

### Utilisation of position-specific nitrogen atoms by *M. tuberculosis*

An essential step in the analysis of nitrogen metabolism with nitrogen sources containing more than one nitrogen atom, such as Gln and Asn, is to define which nitrogen atom(s) is/are being utilised. The ability to utilisation different nitrogen atoms is likely variable and species-specific. To define how *M. tuberculosis* utilise different nitrogen atoms we performed labelling experiments with position-specific labelled Gln and Asn (**Fig. 4a**). The most direct chemical reactions producing 5 key amino acids (namely Glu, Gln, Asp, Asn and Ala) and the label incorporation data obtained from doubly and position-specific labelled ^15^N-Gln and ^15^N-Asn are shown in **Fig. 4b-f.** These results indicate that both nitrogen atoms from Gln and Asn are utilised by *M. tuberculosis* and, specifically that: (i) glutamate synthase is converting the δ-^15^N from Gln into α-^15^N-Glu (**Fig 4b**); this explains the incorporation of δ-15N from Gln into the α-^15^N-Asp, via direct transamination from α-^15^N-Glu (**Fig. 4d**); (ii) direct transamination between α-^15^N-Glu, the other product of the glutamate synthase reaction, and α-^15^N-Asp is clearly observed (**Fig. 4d**); (iii) when position-specific labelled Asn is used, the dominant form of Asp observed in α-^15^N-Asp (**Fig. 4d)**, indicating that the NH_4_^+^ released by asparaginase is likely assimilated to Gln which is distributed broadly in metabolism (**Fig. 4b, 4c and 4f**), but only modestly to Asp (**Fig. 4d**); (iv) labelled Asn is only detectable when Asn is the nitrogen source (**Fig. 4e**), confirming that no Asn synthesis is taking place in *M. tuberculosis*; (v) use of either position-specific labelled Gln or Asn, leads to identical labelling of Glu (**Fig. 4b**), consistent with access of both α and γ/δ nitrogen atoms; (vi) labelling of Gln with position specific Gln and Asn is indistinguishable, demonstrating that all nitrogen derived from Asn is mobilised through Gln (**Fig. 4c**); and (vii) labelling patterns obtained for Ala in the presence of position-specific labelled Gln and Asn are very similar, indicating again that most of the nitrogen derived from Asn is assimilated first into Gln, and then distributed to other metabolites, reflecting the data shown in **Fig. 4c**.

**Figure 4.**
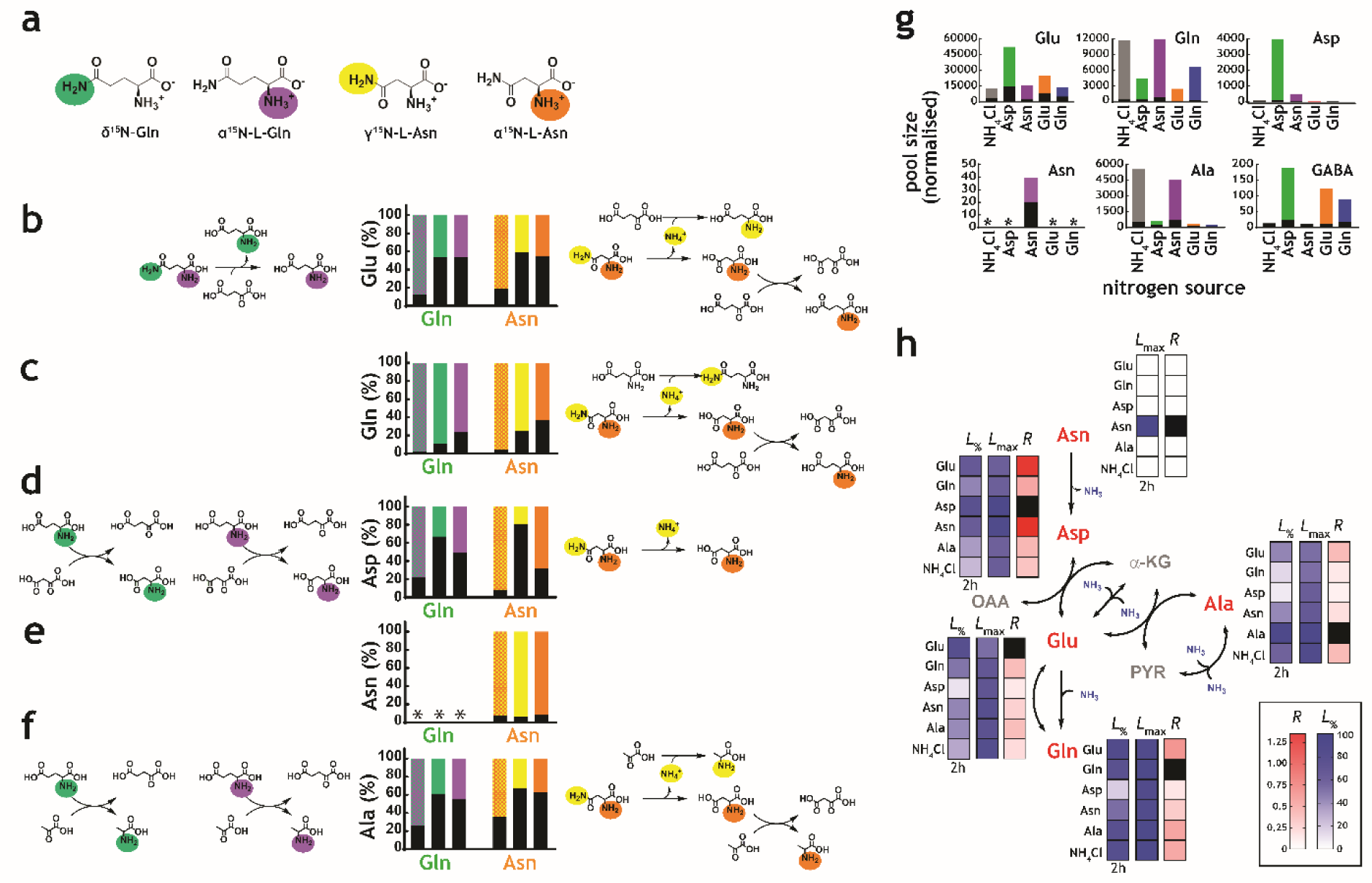
Structural and kinetic analysis of nitrogen utilisation by *M. tuberculosis*. (**a**) Scheme illustrating the structure and position-specific labelling of nitrogen atoms on Gln and Asn. (**b-f**) Data on universally or position-specific labelled Gln or Asn and potential reactions that would lead to the expected labelling patterns obtained. (**b**) Labelling of Glu. Glutamate synthase (with Gln) and asparaginase, glutamate dehydrogenase and glutamate/oxaloacetate transaminase (with Asn). (**c**) Labelling of Gln. Asparaginase, glutamate dehydrogenase, glutamine synthetase (not shown) and glutamate/oxaloacetate transaminase, followed by glutamine synthetase (not shown). (**d**) Labelling of Asp. Glutamate synthase (not shown) and glutamate/oxaloacetate transaminase, with Gln. Asparginase is responsible for most of the labelling in Asp, when Asn is the sole nitrogen source. (**e**) Labelling of Asn. No Asn can be measured in Gln as sole nitrogen source. And most Asn is labelled when Asn is the sole nitrogen source. (**f**) Labelling of Ala. Glutamate synthase (not shown), glutamate/pyruvate transaminase, with Gln as sole nitrogen source. Asparaginase, alanine dehydrogenase and aspartate/pyruvate transaminase, with Asn as sole nitrogen source. (**g**) Representative labelling and pool sizes for different amino acids obtained after 17 h culture in ^15^N-labelled nitrogen sources. (**h**) Data illustrating quantitative analysis of nitrogen labelling in *M. tuberculosis* in sole nitrogen sources. Black squares indicate uptake (cognate amino acid) and not metabolic labelling. Data shown is representative of two independent experiments.

### Kinetics of nitrogen metabolism in *M. tuberculosis*

Label incorporation from ^15^N Glu, ^15^N_2_-Gln, ^15^N-Asp, ^15^N_2_-Asn and ^15^NH_4_Cl obtained under metabolic steady-state, over the course of 17 hours, revealed several important features of *M. tuberculosis* nitrogen metabolism, including different kinetics of ^15^N labelling (**Fig. 4g, 3h, S4**). As expected, regardless of the nitrogen source, robust label incorporation into amino acids belonging to core nitrogen metabolism was observed, with exception of Asn, which was only observed when cells grew in Asn as sole nitrogen source (**Fig. 4g**). It is noteworthy that the Ala pool size and labelling was significantly higher when NH_4_Cl or Asn was the sole nitrogen source. Also, in agreement with data shown in **Fig. 2a**, external amino acid availability does not necessarily correlate with increased intracellular pool size. For example, Glu and Gln are more abundant with Asp and Asn as the sole nitrogen source, respectively (rather than in the cognate amino acid as sole nitrogen source).

Taking the position specific labelling data and corresponding likely metabolic paths, in conjunction with current biochemical and genetic knowledge of the enzymes of the core nitrogen metabolic network (summarised in **Fig. 1**), we calculated exponential labelling rates (*R*) and maximum labelling levels (*L*_max_) for various core amino acids when Asp, Asn, Glu, Gln and NH_4_Cl were used as sole nitrogen sources (**Fig. 4h and Fig. S4**). *L*_max_ for different sole nitrogen sources appears to be similar (**Fig. 4h**), indicating that in principle, nitrogen derived from Glu, Gln, Asp, Asn and NH_4_^+^ can reach similar high levels (close to 100%) in the core nitrogen metabolites of *M. tuberculosis* before the first division (17 h). (**Fig. 4h**). In contrast, *R* values varied considerably, depending on the sole nitrogen source present and reactions needed to transfer the ^15^N atom to individual metabolites (**Fig. 4h and Fig. S4**). Again, NH_4_^+^ is not the most efficient nitrogen source for *M. tuberculosis*, as it leads to only modest labelling of key core metabolites, compared to other sole nitrogen sources. *L*_max_ and *R* for Asn are only consistently observed when Asn is used as sole nitrogen source, supporting the idea that *M. tuberculosis* has a very small Asn pool (**Fig. 4g** and **4h**). *L*_%_ in **Fig. 4h** illustrates early (2 h incubation) labelling of metabolites, and further highlights the differences in *R* values for each nitrogen source. These differences are only related to nitrogen metabolism, as no significant differences are observed in pool sizes of pyruvate, α-ketoglutarate, L-malate and succinate, which are reporting on glycolysis and Krebs cycle metabolism.

### Co-catabolism of amino acids does not improve growth

*M. tuberculosis* has recently been shown to co-catabolise carbon sources simultaneously^21^, leading to better growth than in individual carbon sources. Co-catabolism of carbon sources is a metabolic feature highly unusual in bacteria, which usually catabolise carbon sources sequentially, displaying biphasic (diauxic) growth kinetics. To investigate nitrogen source co-metabolism in *M. tuberculosis*, we grew cells in media containing the following combinations of nitrogen sources: ^15^N-Glu+^14^N-Gln, ^14^N-Glu+U^15^N-Gln, ^15^N-Asp+^14^N-Asn, or ^14^N-Asp+U^15^N-Asn. All nitrogen source combinations lead to robust labelling of Glu, Gln, Asp, Asn (**Fig. 5**), indicating that *M. tuberculosis* is indeed able to take up and co-metabolise nitrogen sources. Extracted ion chromatograms (**Fig. 5a**) illustrate significant ^15^N metabolism in all conditions (*i.e*. high levels of labelled metabolites in the absence of labelled nitrogen sources). Mass spectral data (**Fig. 5b**) reveals that Gln (initially labelled or unlabelled) has been metabolised extensively, generating all three isotopologues (^14^N_2_, ^14^N^15^N and ^15^N_2_). Mass spectral analysis further confirms that no Asn is being synthesised in *M. tuberculosis*, as no labelled Asn (^15^N_2_ or ^15^N_1_) is found when ^15^N-Asp is used as nitrogen source and no ^15^N-Asn is present when ^15^N_2_-Asn is provided as nitrogen source. **Fig. 5c** provides average values and errors for labelling of Glu, Gln, Asp, Asn and Ala, in dual nitrogen sources. Interestingly, Ala labelling appears to derive mainly from Asn, rather than Asp. This suggests that Asn is being hydrolysed to Asp and NH_4_^+^, and likely that Ala is either a main entry point for ^15^NH_4_^+^ or it serves as nitrogen storage. This result is in strict agreement with the very fast and extensive label incorporation of Ala when using ^15^NH_4_^+^ or ^15^N-Asn as sole nitrogen sources (**Fig. 4g** and **4h**). In spite of clear co-metabolism of two different nitrogen sources (*i.e.* Glu/Gln and Asp/Asn), no growth advantage (faster doubling time and highest biomass) is observed (**Fig. S6**).

**Figure 5.**
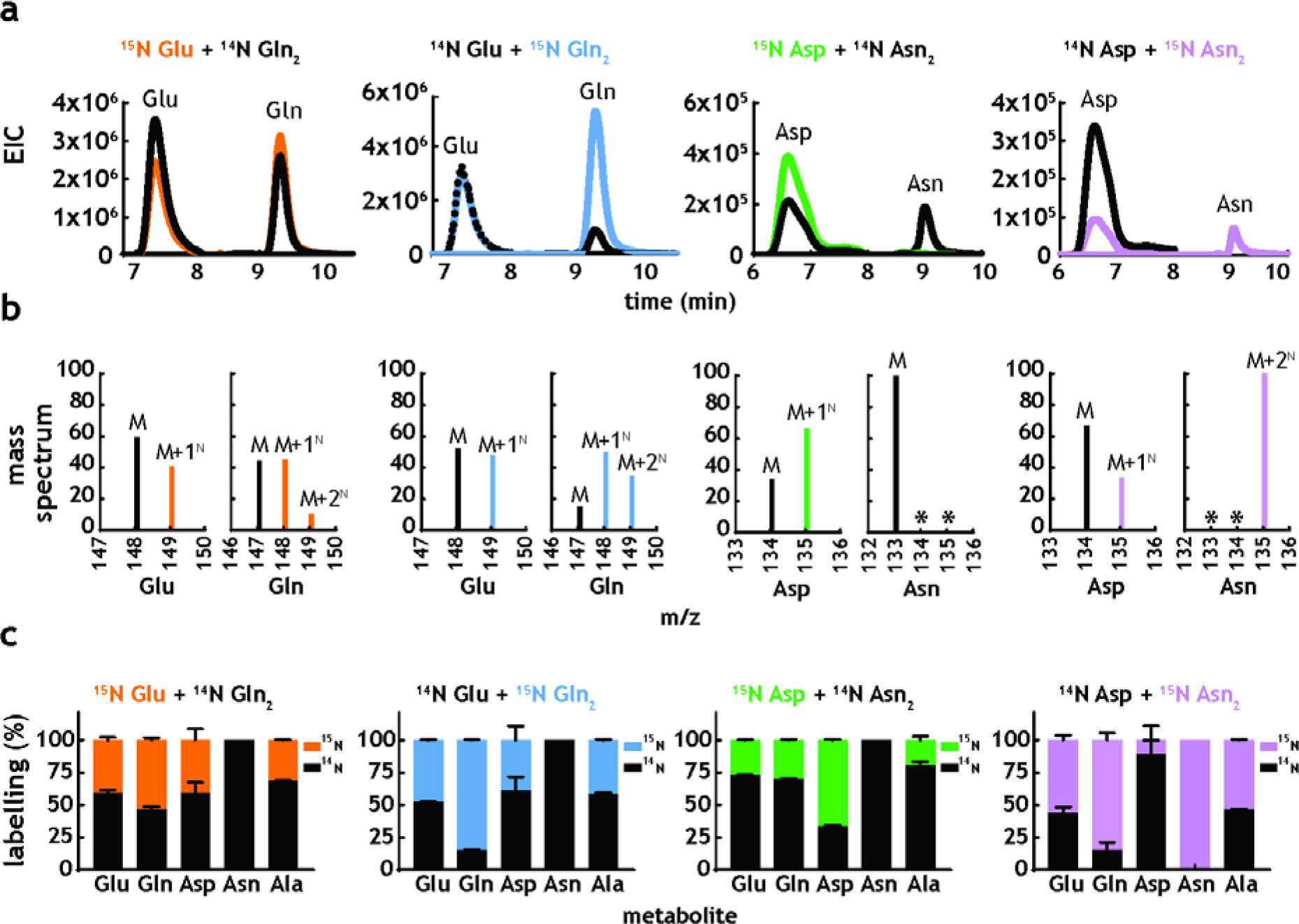
*M. tuberculosis* co-metabolises nitrogen sources. (**a**) Representative extracted ion chromatograms (EICs) for intracellular metabolites from cultures obtained in the presence of Glu+Gln or Asp+Asn, with one of the nitrogen sources ^15^N-labelled. (**b**) Representative mass spectra corresponding to the metabolites in **Fig. 5a**. (**c**) Combined labelling data obtained for the same metabolites, in different combinations of two carbon sources. Bars are averages of three biological replicates and error bars are the standard error of the mean.

### Alanine and alanine dehydrogenase as a fundamental node in nitrogen metabolism

Ala pool size and labelling patterns (**Fig. 4g** and **4h**) are incompatible with our current understanding of nitrogen metabolism in *M. tuberculosis*. If NH_4_^+^ utilization, either direct or derived from Asn, proceeded through glutamine synthetase or glutamate dehydrogenase, labelling of Glu would always be greater than Ala, which would be produced by transamination of Glu. However, this is not the case. To investigate Ala metabolism in the context of nitrogen assimilation, we first confirmed whether Ala could serve as a nitrogen source. **Fig. 6a** shows that *M. tuberculosis* can grow in the presence of Ala as a sole nitrogen source, or in binary combination of Ala with Glu, Gln, Asp, Asn, or NH_4_^+^Cl. These results are consistent with Ala being utilised as a sole nitrogen source and in combination with other nitrogen sources, but without any growth advantage (mirroring the result observed for Glu/Gln and Asp/Asn co-metabolism). qPCR analysis of transcript levels for asparaginase (*ansA*), glutamine synthetase (*glnA1*), glutamate dehydrogenase (*gdh*) and alanine dehydrogenase (*ald*), in sole nitrogen sources was carried out to define if transcriptional programmes are involved in control of amino acid metabolism, and in particular of alanine dehydrogenase, despite the current lack of potential transcriptional regulators of nitrogen metabolism (**Fig. 6b**). Consistent the hypothesis that alanine dehydrogenase works as a NH_4_^+^ assimilatory route, *ald* RNA levels are found to be higher when *M. tuberculosis* was grown in media with NH_4_^+^, Asn, Asp and Gln, compared to nitrogen free medium (**Fig. 6b, -N/N+**). In addition, *ald* RNA levels are found to be decreased under nitrogen starved conditions, in comparison to *gdh*, *glnA1* and *ansA* RNA levels, suggesting that *ald*-driven nitrogen assimilation is likely more important under nitrogen-rich conditions.

**Figure 6.**
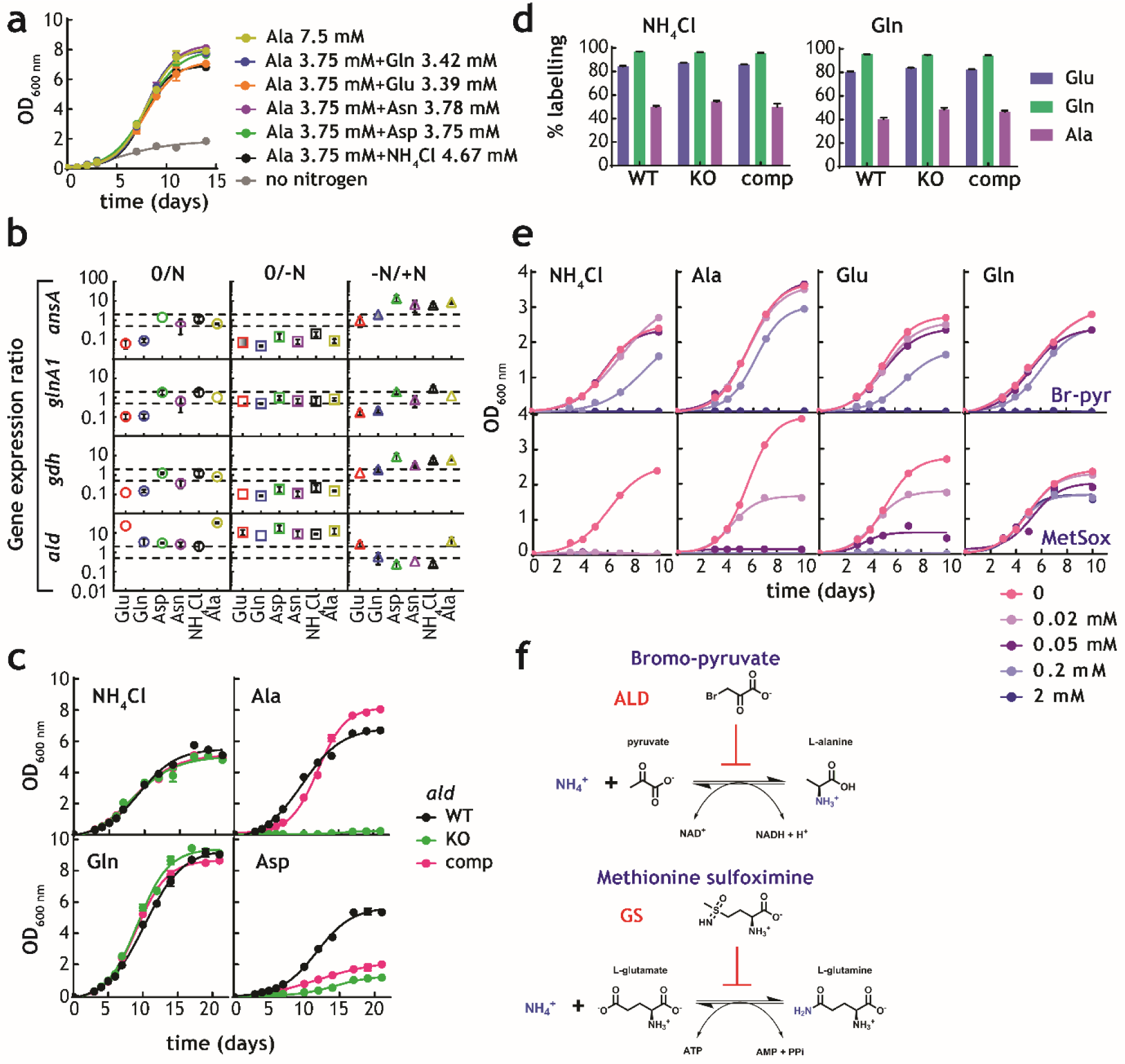
Alanine and alanine dehydrogenase roles in *M. tuberculosis* nitrogen metabolism. (**a**) Growth of *M. tuberculosis* on alanine as sole nitrogen source or in combination with a second nitrogen source. (**b**) Gene expression ratios in different nitrogen sources. (**c**) Parent, *ald* KO, and complemented strains growth in selected sole nitrogen sources. (**d**) Labelling of selected amino acids obtained with parent, *ald* KO, and complemented strains cultured in NH_4_^+^ or Gln as sole nitrogen sources. (**e**) Growth of *M. tuberculosis* in single nitrogen sources in the presence of various concentrations of bromo-pyruvate or methionine sulfoximine, inhibitors of alanine dehydrogenase and glutamine synthethase, respectively. (**f**) Reaction catalysed by alanine dehydrogenase and glutamine synthetase and their inhibitors. Data shown are representative of two independent experiments. Error bars are standard error of the mean.

To define the role of *ald*-encoded alanine dehydrogenase in mobilisation of nitrogen to and from Ala, we compared a *M. tuberculosis* lacking *ald*, to parent and complemented strains ^23^. Genetic disruption of *ald* abolished the ability of *M. tuberculosis* to grow when Ala was the sole nitrogen source, with no effect when NH_4_^+^ or Gln were sole nitrogen sources (**Fig. 6c**). These results demonstrate that alanine dehydrogenase is essential for assimilation of NH_4_^+^ from Ala, as shown elsewhere ^23^. Interestingly, growth was also significantly diminished in the *ald*-knockout strain when Asp was the sole nitrogen soure, however only partial complementation was obtained (**Fig. 6c**). Growth of parent, *ald*-knockout and complemented strains was indistinguishable in NH_4_^+^ as sole nitrogen source, confirming that *ald* is not the main route for NH_4_^+^ assimilation in *M. tuberculosis*. This secondary role of alanine dehydrogenase in assimilation is further supported by lack of changes in label incorporation into Glu, Gln and Ala when parent, *ald*-knockout, and complemented strains were grown with ^15^NH_4_^+^Cl or ^15^N-Gln (**Fig. 6d**). Final evidence for the essentiality of alanine dehydrogenase in nitrogen assimilation from Ala and secondary role during ^15^N assimilation was obtained using the inhibitors of alanine dehydrogenase and glutamine synthetase, bromo-pyruvate^24^ and methionine sulfoximine^25^, respectively (**Fig. 6e, 6f**). Bromo-pyruvate partially inhibits growth when NH_4_^+^, Ala and Glu are used as sole nitrogen sources, but not in Gln. In addition to alanine dehydrogenase, glutamate dehydrogenase is likely also partially inhibited at the concentrations tested, leading to the phenotype observed in Glu (**Fig. 6f**, top panels). Methionine sulfoximine abrogated growth in NH_4_^+^ and to a less extent in Ala and Glu, but not in Gln as the sole nitrogen sources (**Fig. 6f**, bottom panels). Together, these results demonstrate that alanine dehydrogenase is essential for utilisation of Ala as sole nitrogen source and secondary to the utilisation of NH_4_^+^, a task undertaken primarily by glutamine synthetase.

## Discussion

*M. tuberculosis* is a metabolic generalist, that is, it can make the molecules it requires when provided with simple carbon and nitrogen sources. Despite that, *M. tuberculosis* confinement to the human host over several thousand years has dramatically altered its ability to metabolise host-derived nutrients, which are not necessarily abundant in the environment. In this work, we have uncovered several traits that lead to flexible utilisation of amino acids as nitrogen sources. Most strikingly, *M. tuberculosis* appears to not tightly control amino acid pool sizes; be able to co-metabolise amino acids as nitrogen sources; and employ alanine dehydrogenase as an assimilatory route for NH_4_^+^. In sharp contrast to most non-nitrogen fixing bacteria studied to date, *M. tuberculosis* appears to prefer to utilise amino acids such as Glu, Gln, Asp, Asn and Ala as nitrogen sources, instead of NH_4_^+^.

## Materials and methods

### Strains and growth media

*M. tuberculosis* H37Rv was used for growth and metabolic studies. Alanine dehydrogenase Rv2780 knockout, parent (*M. tuberculosis* H37Rv), and complemented strains were generated previously^23^. Liquid media used for *M. tuberculosis* growth: (a) commercially available Middlebrook 7H9 (Sigma UK) supplemented with (wt/vol) 0.05% tyloxapol, (wt/vol) 0.4% glycerol and albumin-dextrose-catalase (ADC) supplement (Sigma) (b) synthetic 7H9Nx (0.5 g/L sodium sulphate, 0.1 g/L sodium citrate, 1 mg/L pyridoxine, 0.5 mg/L biotin, 2.5 g/L sodium phosphate dibasic, 1.0 g/L monobasic potassium phosphate, 0.04 g/L ferric ammonium citrate, 0.05 g/L magnesium sulphate, 0.5 mg/L calcium chloride, 1 mg/L zinc sulphate, 1 mg/L copper sulphate, 0.4% glycerol, 0.05% tyloxapol, 10% ADC, pH to 6.6) and supplemented with nitrogen source of interest **(c)** commercially available Middlebrook 7H10 (Sigma UK) supplemented with 0.5% glycerol and (vol/vol) oleic acid-albumin-dextrose-catalase (OADC) supplement (d) synthetic 7H10Nx (sodium citrate 0.4 g/L, copper sulfate 1 mg/L, calcium chloride 0.5 mg/L, zinc sulphate 1 mg/L, magnesium sulphate 0.025 g/L, ferric ammonium citrate 0.04 g/L, malachite green 0.25 mg/L, biotin 0.5 mg/L, pyridoxine hydrochloride 1 mg/L, sodium sulfate 0.5 g/L, monopotassium phosphate 1.5 g/L, disodium phosphate 1.5 g/L, Agar 15 g/L, 0.5% glycerol) and supplemented with 10% OADC.

### Metabolite extraction

*M. tuberculosis* was grown in liquid media to mid logarithmic phase and then 1 ml of culture was transferred on 0.22 µm nitrocellulose filter (GSWP02500, Millipore) using vacuum filtration and placed on 7H10Nx agar plates. *M. tuberculoisis* loaded filters were then grown at 37˚C for 5 days. On day 5 filters were transferred on chemically identical ^15^N 7H10Nx plates for isotopic labelling and metabolites were extracted with acetonitrile/methanol/_d_H2O 2:2:1 (v/v/v) at -40°C. Cells were then mechanically disrupted using a Fastprep ryboliser (QBiogene). Samples were centrifuged for 10 min 13,000 rpm at 4˚C, and the supernatant was recovered and filtered through 0.22µm spin-X centrifuge tube filter (8160, Costar).

### Liquid chromatography-mass spectrometry (LC-MS)

A Cogent Diamond Hydride Type C column (Agilent) was used for normal phase chromatography on 1200 LC system (Agilent) coupled to an Accurate Mass 6220 TOF (Agilent) mass spectrometer fitted with an MultiMode ion source. Metabolite extracts were mixed with solvent A 1:1 and separated using mobile phase of solvent gradient A and B: 0–2 min, 85% B; 3–5 min, 80% B; 6–7 min, 75%; 8–9 min, 70% B; 10–11.1 min, 50% B; 11.1–14:10 min 20% B; 14:10-18:10 5% B; 18:10-19 85% B. Solvent A was acetonitrile with 0.2% acetic acid and solvent B was ddH2O with 0.2 % acetic acid. Reference mass solution (G1969-85001, Agilent Technologies) was used for continuous mass axis calibration. Analytical amino acids standard mix (Fluka A9906) was used for retention time match. Ions were identified based on their accurate mass, retention time and spectral information, yielding errors below 5 ppm. Spectra were analysed using MassHunter Qualitative analysis B.07.00 and MassHunter Profinder B.08.00 software. Statistical validation of samples/runs were performed using principal component analysis, using Mass Profiler Professional (B.07.01).

### Extraction and analysis of RNA, and qPCR

*M. tuberculosis* was pre-adapted in 7H9Nx medium for 3 days and then grown in identical medium. Cells were harvested at an OD_600_ between 0.8 and 1.0. RNA was extracted using Fast RNA Pro Blue kit according to manufacturer’s instructions. DNA was removed by treatment with 3 U RNase-free DNase using the TURBO DNA-free kit (Ambion) according to the manufacturer’s instructions and cleaned following RNeasy Mini kit (Qiagen). The concentration of the RNA was determined using a NanoDrop One (Thermo) (Promega) spectrophotometer. Reverse transcriptase PCR was performed using SuperScript IV (Invitrogen), according to the manufacturer’s instructions for cDNA synthesis. After cDNA synthesis, qPCR was carried out using the PowerUp SYBR Green Master Mix with ROX (Applied Biosystems) on a QuantStudio 7 Flex Real-Time PCR System. SigE (Rv1221) was used as an internal standard, and the ddCt method was used for the calculation of gene expression ratios. Error bars represent standard deviations from three biological replicates.

## Acknowledgements

We thank Dr James McRae for critical reading of the manuscript. This work was primarily supported by a Wellcome Trust New Investigator Award (104785/B/14/Z) to L.P.S.C. The L.P.S.C. lab is also funded by the Francis Crick Institute, which receives its core funding from Cancer Research UK (FC001060), the UK Medical Research Council (FC001060), and the Wellcome Trust (FC001060).

## Author contributions

AA, DMH, MP, AGG, carried out experiments; AA, MP, LPSC analysed data; AA, MP, CDS, LPSC interpreted data; AA, MP, LPSC prepared figures; AA, LPSC wrote and revised the manuscript.

## References

1. WHO. Global Tuberculosis Report. (WHO, Geneve, 2017).

2. Barry, C. E., 3rd et al. The spectrum of latent tuberculosis: rethinking the biology and intervention strategies. Nat Rev Microbiol 7, 845–855, doi:10.1038/nrmicro2236 (2009).

3. Ehrt, S., Schnappinger, D. & Rhee, K. Y. Metabolic principles of persistence and pathogenicity in Mycobacterium tuberculosis. Nat Rev Microbiol, doi:10.1038/s41579-018-0013-4 (2018).

4. Rhee, K. Y. et al. Central carbon metabolism in Mycobacterium tuberculosis: an unexpected frontier. Trends Microbiol 19, 307–314, doi:10.1016/j.tim.2011.03.008 (2011).

5. Gouzy, A., Poquet, Y. & Neyrolles, O. Nitrogen metabolism in Mycobacterium tuberculosis physiology and virulence. Nat Rev Microbiol 12, 729–737, doi:10.1038/nrmicro3349 (2014).

6. Carroll, P., Pashley, C. A. & Parish, T. Functional analysis of GlnE, an essential adenylyl transferase in Mycobacterium tuberculosis. J Bacteriol 190, 4894–4902, doi:10.1128/JB.00166-08 (2008).

7. Cowley, S. et al. The Mycobacterium tuberculosis protein serine/threonine kinase PknG is linked to cellular glutamate/glutamine levels and is important for growth in vivo. Mol Microbiol 52, 1691–1702, doi:10.1111/j.1365-2958.2004.04085.x (2004).

8. Nott, T. J. et al. An intramolecular switch regulates phosphoindependent FHA domain interactions in Mycobacterium tuberculosis. Sci Signal 2, ra12, doi:10.1126/scisignal.2000212 (2009).

9. O’Hare, H. M. et al. Regulation of glutamate metabolism by protein kinases in mycobacteria. Mol Microbiol 70, 1408–1423, doi:10.1111/j.1365-2958.2008.06489.x (2008).

10. Pashley, C. A., Brown, A. C., Robertson, D. & Parish, T. Identification of the Mycobacterium tuberculosis GlnE promoter and its response to nitrogen availability. Microbiology 152, 2727–2734, doi:10.1099/mic.0.28942-0 (2006).

11. Read, R., Pashley, C. A., Smith, D. & Parish, T. The role of GlnD in ammonia assimilation in Mycobacterium tuberculosis. Tuberculosis (Edinb) 87, 384–390, doi:10.1016/j.tube.2006.12.003 (2007).

12. Rieck, B. et al. PknG senses amino acid availability to control metabolism and virulence of Mycobacterium tuberculosis. PLoS Pathog 13, e1006399, doi:10.1371/journal.ppat.1006399 (2017).

13. Parish, T. & Stoker, N. G. glnE is an essential gene in Mycobacterium tuberculosis. J Bacteriol 182, 5715–5720 (2000).

14. Williams, K. J. et al. Deciphering the metabolic response of Mycobacterium tuberculosis to nitrogen stress. Mol Microbiol 97, 1142–1157, doi:10.1111/mmi.13091 (2015).

15. Liu, X. X., Shen, M. J., Liu, W. B. & Ye, B. C. GlnR-Mediated Regulation of Short-Chain Fatty Acid Assimilation in Mycobacterium smegmatis. Front Microbiol 9, 1311, doi:10.3389/fmicb.2018.01311 (2018).

16. Petridis, M., Benjak, A. & Cook, G. M. Defining the nitrogen regulated transcriptome of Mycobacterium smegmatis using continuous culture. BMC Genomics 16, 821, doi:10.1186/s12864-015-2051-x (2015).

17. Gouzy, A. et al. Mycobacterium tuberculosis exploits asparagine to assimilate nitrogen and resist acid stress during infection. PLoS Pathog 10, e1003928, doi:10.1371/journal.ppat.1003928 (2014).

18. Gouzy, A. et al. Mycobacterium tuberculosis nitrogen assimilation and host colonization require aspartate. Nat Chem Biol 9, 674–676, doi:10.1038/nchembio.1355 (2013).

19. Berney, M. et al. Essential roles of methionine and S-adenosylmethionine in the autarkic lifestyle of Mycobacterium tuberculosis. Proc Natl Acad Sci U S A 112, 10008–10013, doi:10.1073/pnas.1513033112 (2015).

20. Hondalus, M. K. et al. Attenuation of and protection induced by a leucine auxotroph of Mycobacterium tuberculosis. Infect Immun 68, 2888–2898 (2000).

21. de Carvalho, L. P. et al. Metabolomics of Mycobacterium tuberculosis reveals compartmentalized co-catabolism of carbon substrates. Chem Biol 17, 1122–1131, doi:10.1016/j.chembiol.2010.08.009 (2010).

22. Larrouy-Maumus, G. et al. Discovery of a glycerol 3-phosphate phosphatase reveals glycerophospholipid polar head recycling in Mycobacterium tuberculosis. Proc Natl Acad Sci U S A 110, 11320–11325, doi:10.1073/pnas.1221597110 (2013).

23. Giffin, M. M., Modesti, L., Raab, R. W., Wayne, L. G. & Sohaskey, C. D. ald of Mycobacterium tuberculosis encodes both the alanine dehydrogenase and the putative glycine dehydrogenase. J Bacteriol 194, 1045–1054, doi:10.1128/JB.05914-11 (2012).

24. Bellion, E. & Tan, F. An NAD+-dependent alanine dehydrogenase from a methylotrophic bacterium. Biochem J 244, 565–570 (1987).

25. Harth, G. & Horwitz, M. A. An inhibitor of exported Mycobacterium tuberculosis glutamine synthetase selectively blocks the growth of pathogenic mycobacteria in axenic culture and in human monocytes: extracellular proteins as potential novel drug targets. J Exp Med 189, 1425–1436 (1999).

